# Eukaryotic Lagging Strand Synthesis is Distributive

**DOI:** 10.1101/2025.09.17.676973

**Authors:** Luke Lynch, Gheorghe Chistol

## Abstract

DNA’s anti-parallel structure presents a topological challenge to the replisome, as the leading and lagging strands must be synthesized in opposite directions. It is widely believed that the lagging strand polymerases Pol δ and Pol α are stably tethered to the eukaryotic replisome to facilitate their recycling and to coordinate leading and lagging strand synthesis. To test this idea, we directly visualized the dynamics of Pol α/δ at active replication forks via single-molecule imaging in *Xenopus* nuclear extracts. Surprisingly, we find that neither Pol α nor Pol δ is stably tethered to the eukaryotic replisome. Instead, lagging strand synthesis occurs *distributively* and entails the recruitment of new molecules of Pol α and Pol δ to facilitate the synthesis of new Okazaki fragments. Our data reveal a highly dynamic mechanism of lagging strand synthesis and challenge the current textbook model of the eukaryotic replisome.

## INTRODUCTION

Organisms with double-stranded DNA genomes universally perform continuous leading strand synthesis and discontinuous lagging strand synthesis. In the eukaryotic replisome, Polymerase ε(Pol ε) synthesizes the leading strand continuously, while Polymerase α-Primase (Pol α, also referred to as POLA) and Polymerase δ (Pol δ, also known as POLD) synthesize the lagging strand discontinuously^1–4^.

During lagging strand synthesis, Pol α is recruited to the replicative DNA helicase CMG (CDC45•MCM2-7•GINS) and engages the lagging strand template^5^. First, Pol α’s primase subunits (PRIM1 and PRIM2) synthesize a 6-10 ribonucleotide (rNTP) primer^6^. Second, Pol α’s polymerase subunits (POLA1 and POLA2) extend the RNA primer by 20-30 deoxyribonucleotides (dNTPs), then disengage the template DNA^7,8^. Next, the Proliferating Cell Nuclear Antigen processivity clamp (PCNA) is loaded onto the primer-template junction, promoting the recruitment of Pol δ^9^. Pol δ extends the primer by several hundred nucleotides^10^ until colliding with a downstream Okazaki fragment^11^. Finally, the Flap Endo-Nuclease 1 (FEN1) cleaves the short RNA primer from the 5’ end of the Okazaki fragment and neighboring lagging strand fragments are ligated^10^.

DNA’s anti-parallel structure presents a topological challenge to the replisome, as the leading and lagging strands must be synthesized in opposite directions. Early studies of bacteriophage replication revealed that both the DNA Primase and the lagging strand DNA polymerase are stably tethered to the bacteriophage replisome, enabling it to processively synthesize many consecutive Okazaki fragments. To rationalize these observations, it was proposed that the lagging strand DNA template loops to facilitate the coordinated synthesis of leading and lagging strands by a single stable enzyme complex^12,13^. This was coined as the “trombone” model, referring to the trombone-like loop formed by the lagging strand template.

While originally proposed for bacteriophage replication, the trombone model is widely accepted as the textbook model of eukaryotic DNA replication^14,15^ (Figure 1A). It is a very attractive model because it provides a way to coordinate the synthesis of nascent leading and lagging strands. This coordination is thought to be necessary for preventing ssDNA accumulation and ensuring genome stability^15,16^.

**Figure 1:**
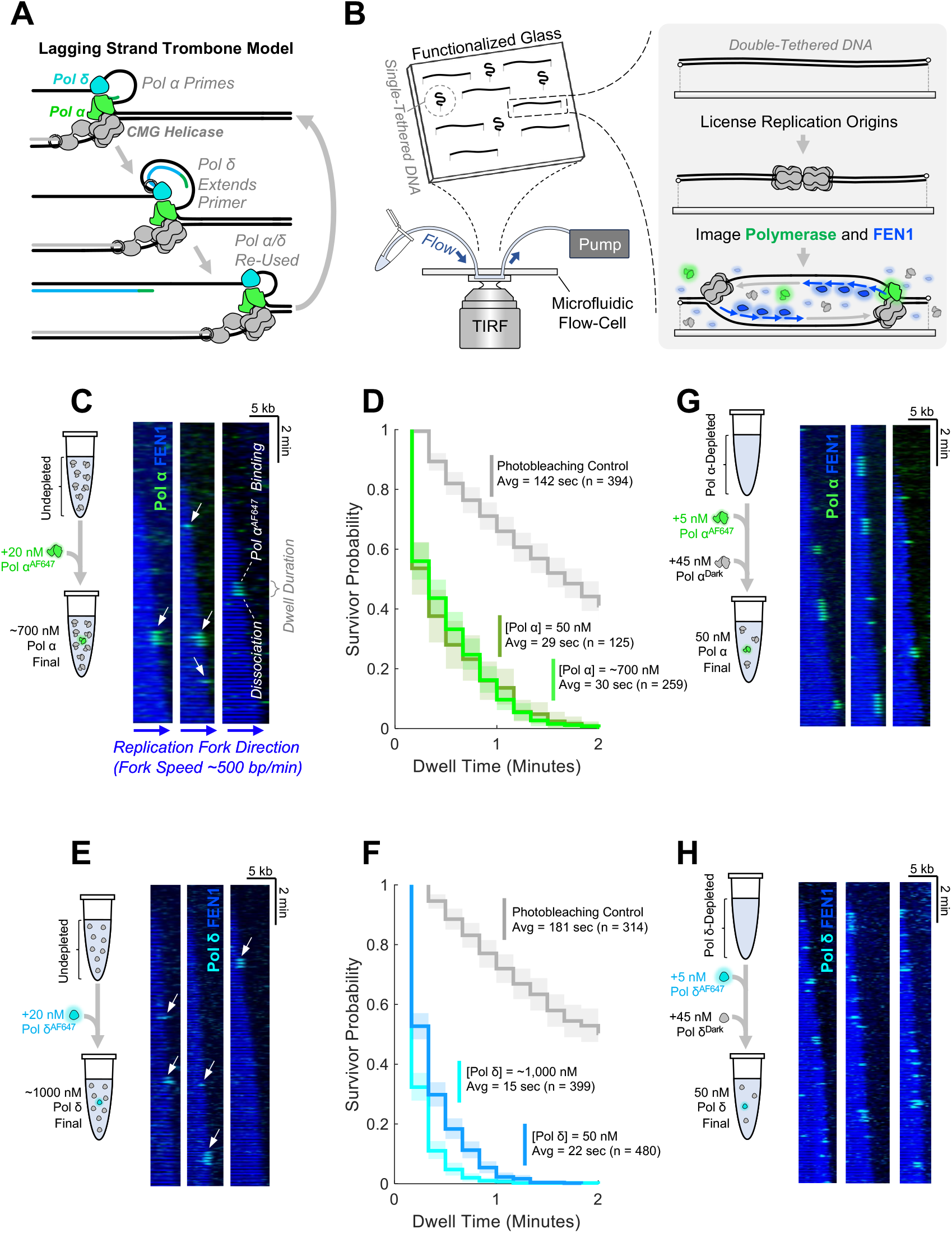
Pol α/δ Are Not Stably Tethered to the Replisom. **(A)**Illustration of the trombone model with continuous recycling of Pol α/δ molecules tethered to the replisome.**(B)**Diagram of our single-molecule imaging workflow. **(C)** Representative kymograms of Pol α^AF647^ (green) and FEN1^mKikGR^ (blue) from an experiment where 20 nM of recombinant Pol α^AF647^ was spiked in to undepleted extract containing ∼700 nM of endogenous Pol α. **(D)** Survivor probability curves of Pol α^AF647^ binding event dwell times. The Pol α^AF647^ photobleaching control is shown in gray. **(E)** Representative kymograms of Pol δ^AF647^ (cyan) and FEN1^mKikGR^ (blue) from an experiment where 20 nM of Pol δ^AF647^ was spiked in to undepleted extract containing ∼1,000 nM of endogenous Pol δ. **(F)** Survivor probability curves of Pol δ^AF647^ binding event dwell times. The Pol δ^AF647^ photobleaching control is shown in gray. **(G)** Representative kymograms of Pol α^AF647^ (green) and FEN1^mKikGR^ (blue) from an experiment where 5 nM Pol α^AF647^ was spiked into Pol α-depleted extract supplemented with 45 nM unlabeled Pol α. **(H)** Representative kymograms of Pol δ^AF647^ (green) and FEN1^mKikGR^ (blue) from an experiment where 5 nM Pol δ^AF647^ was spiked into Pol δ-depleted extract supplemented with 45 nM unlabeled Pol δ. In panels **(D)** and **(F)**, semi-transparent boxes depict the 95% Confidence Interval (CI) estimated via bootstrapping.

The trombone model for eukaryotic DNA replication is strongly supported by two recent yeast studies. A minimal *in vitro* reconstitution study reported that Pol α and Pol δ travel with the replisome for several minutes – long enough to synthesize hundreds of Okazaki fragments^17^. Another study imaged the diffusion of Pol α and Pol δ molecules *in vivo* and concluded that lagging strand polymerases are stably recycled for multiple rounds of Okazaki fragment synthesis^18^. Taken together, these observations are consistent with the core premise of the trombone model that Pol α/δ are stably tethered to the eukaryotic replisome (Figure 1A).

To understand how Pol α and Pol δ are tethered to the eukaryotic replisome, and to elucidate how they carry out discontinuous lagging strand synthesis, we used real-time single-molecule imaging to directly visualize Pol α and Pol δ at active replication forks in *Xenopus* nuclear egg extract. This extract contains the entire soluble nuclear proteome and faithfully recapitulates DNA replication and other genome maintenance pathways^19,20^.

Surprisingly, we find that Pol α and Pol δ are not stably tethered to active replisomes. Instead, Pol α and Pol δ are transiently recruited to elongating forks, where they reside for a very short time (15-30 sec) before dissociating. Interestingly, we do observe instances when Pol α resides on DNA for several minutes, but this only occurs at stalled replication forks.

The trombone model predicts that Pol α and Pol δ should be stably tethered to the replisome, and therefore the replisome should resist a chase with catalytically inactive Pol α/δ. Instead, we find that catalytically-dead Pol α/δ rapidly replace wildtype enzymes at the replication fork. This polymerase exchange occurs within ∼40 seconds, i.e. roughly the time needed to synthesize one or two Okazaki fragments. Our experiments also reveal that catalytically-inactive Pol δ molecules are deposited behind advancing replication forks, further supporting the idea that Pol δ is not stably tethered to the replisome. Finally, we show that blocking lagging strand synthesis does not impair CMG progression, indicating that leading and lagging strand synthesis are not intrinsically coupled in eukaryotes.

Taken together, our findings are inconsistent with the classic trombone model of lagging strand synthesis. Instead, our data support a *distributive* mechanism, where new copies of Pol α and Pol δ are constantly recruited to replication forks to sustain the synthesis of new Okazaki fragments.

## RESULTS

### Neither Pol α Nor Pol δ Are Stably Tethered to the Replisome During Unperturbed DNA Replication

The central premise of the trombone model is that lagging strand polymerases are stably tethered to the replisome and therefore should processively travel with active replication forks for several minutes (Figure 1A). To directly visualize this process, we fluorescently labeled recombinant Pol α and Pol δ with the organic dye Alexa-Fluor 647 (Figure S1A-D) and monitored the dynamics of Pol α^AF647^ and Pol δ^AF647^ at replication forks via single-molecule imaging (Figure 1B). An ensemble plasmid replication assay confirmed that fluorescent labeling did not impair the biochemical activity of Pol α/ δ (Figure S1A, S1C).

In our single-molecule assay, linear λ DNA templates were flow-stretched and tethered to a functionalized coverslip in a microfluidic flow-cell (Figure 1B). The flow-cell was incubated with *Xenopus* egg extract which robustly supports origin firing and replication elongation^21^. The reaction was supplemented with recombinant Flap Endonuclease 1 fused to the fluorescent protein mKikGR (FEN1^mKikGR^), which binds nascent Okazaki fragments and serves as a marker of lagging strand synthesis^22^.

First, we spiked 20 nM of Pol α^AF647^ into undepleted extract, which contains ∼700 nM of endogenous Pol α. This enabled us to unambiguously monitor individual polymerase binding and dissociation events. Pol α^AF647^ bound to active replication forks and resided at the fork for an average of 30 seconds before dissociating (Figure 1C, Figure 1D light green curve). 30 seconds corresponds to forks travelling 200-250 nucleotides in our experiments (where the average fork speed is 400-500 nt/min). This is consistent with the length of one or two Okazaki fragments in *Xenopus* egg extract^23^. Although we did observe some long-lived Pol α^AF647^ binding events, they were exclusively associated with stalled forks and therefore do not reflect polymerase dynamics during active replication elongation (Figure S1E).

Second, we spiked 20 nM of Pol δ^AF647^ into undepleted extract, which contains ∼1000 nM of endogenous Pol δ. We found that Pol δ^AF647^ transiently bound to active replication forks with an average residence time of only ∼15 seconds (Figure 1E, Figure 1F light blue curve). Within 15 seconds a replication fork in our experiment travels 100-125 nucleotides, consistent with the size of one Okazaki fragment. Unlike Pol α, Pol δ was not retained at stalled replication forks.

The concentrations of endogenous Pol α (∼700 nM) and Pol δ (∼1000 nM) in undepleted extract are in the same ballpark as the estimated concentrations of Pol α (360-630 nM) and Pol δ (300-650 nM) in human cells^24^ (upper/lower bounds are defined by the least/most abundant subunits, concentrations were corrected for nuclear volume). Since Pol α/δ abundance may vary between cell types, we carried out an ensemble assay to determine the lowest polymerase concentration that supports normal DNA replication (Figure S1F-M). We found that Pol α/δ concentrations above ∼50 nM supported DNA replication as well as undepleted extract (Figure S1H, S1K). Interestingly, polymerase concentrations below 50 nM resulted in elevated pCHK1 levels relative to mock-depleted controls (Figure S1J, S1M).

A recent study using an *in vitro* reconstitution with yeast proteins reported that high Pol α/δ concentrations promote the dissociation of replisome-bound Pol α/δ^9^. Considering this report, we repeated the polymerase spike-in experiment using Pol α-depleted extract supplemented with 50 nM of recombinant Pol α: 5 nM of Pol α^AF647^ and 45 nM of unlabeled Pol α. We used 50 nM as it is the lowest polymerase concentration that supports normal replication. Importantly, lowering the total concentration of Pol α to 50 nM did not significantly change its residence time at the fork (Figure 1G, Figure 1D dark green curve). We conclude that Pol α’s short residence time at the fork is an intrinsic property of this enzyme and does not depend on Pol α’s concentration in extract.

We performed a similar experiment in Pol δ-depleted extract supplemented with 5 nM of Pol δ^AF647^ and 45 nM of unlabeled Pol δ. Pol δ^AF647^ bound to replication forks with an average residence time of only ∼22 sec (Figure 1H, Figure 1F dark blue curve). Interestingly, Pol δ’s residence time is different from that of Pol α, suggesting that they do not form a stable complex as hypothesized previously^17,25,26^. Consistent with this, Pol α and Pol δ co-IP very weakly (Figure S1N). Taken together, these data indicate that neither Pol α nor Pol δ reside at the fork long enough to synthesize several consecutive Okazaki fragments.

### Binding of a Single Copy of Inactive Pol α/δ to the Fork Does Not Impair Lagging Strand Synthesis

The trombone model predicts that the incorporation of a single copy of catalytically inactive Pol α or Pol δ into the replisome should block lagging strand synthesis for several minutes. To test this prediction, we generated a catalytically dead Pol α (Pol α^Dead^) by introducing two point-mutations in the DNA polymerase subunit POLA1 (D998S, D1000S) as described before^17^ (Figure 2A). Notably, Pol α^Dead^ retains its RNA primase activity^17^. We verified that Pol α^Dead^ cannot rescue DNA replication in a Pol α-depleted extract (Figure 2B, S2A). Spiking 20 nM of fluorescently labeled Pol α^Dead^ into undepleted extract revealed that it binds replication forks and resides on DNA for a similar amount of time as wildtype Pol α (Figure 2C-D). Importantly, the recruitment of Pol α^Dead^ to the replisome did not impair lagging strand synthesis as monitored by the growth of the FEN1^mKikGR^ tract (Figure 2C).

**Figure 2:**
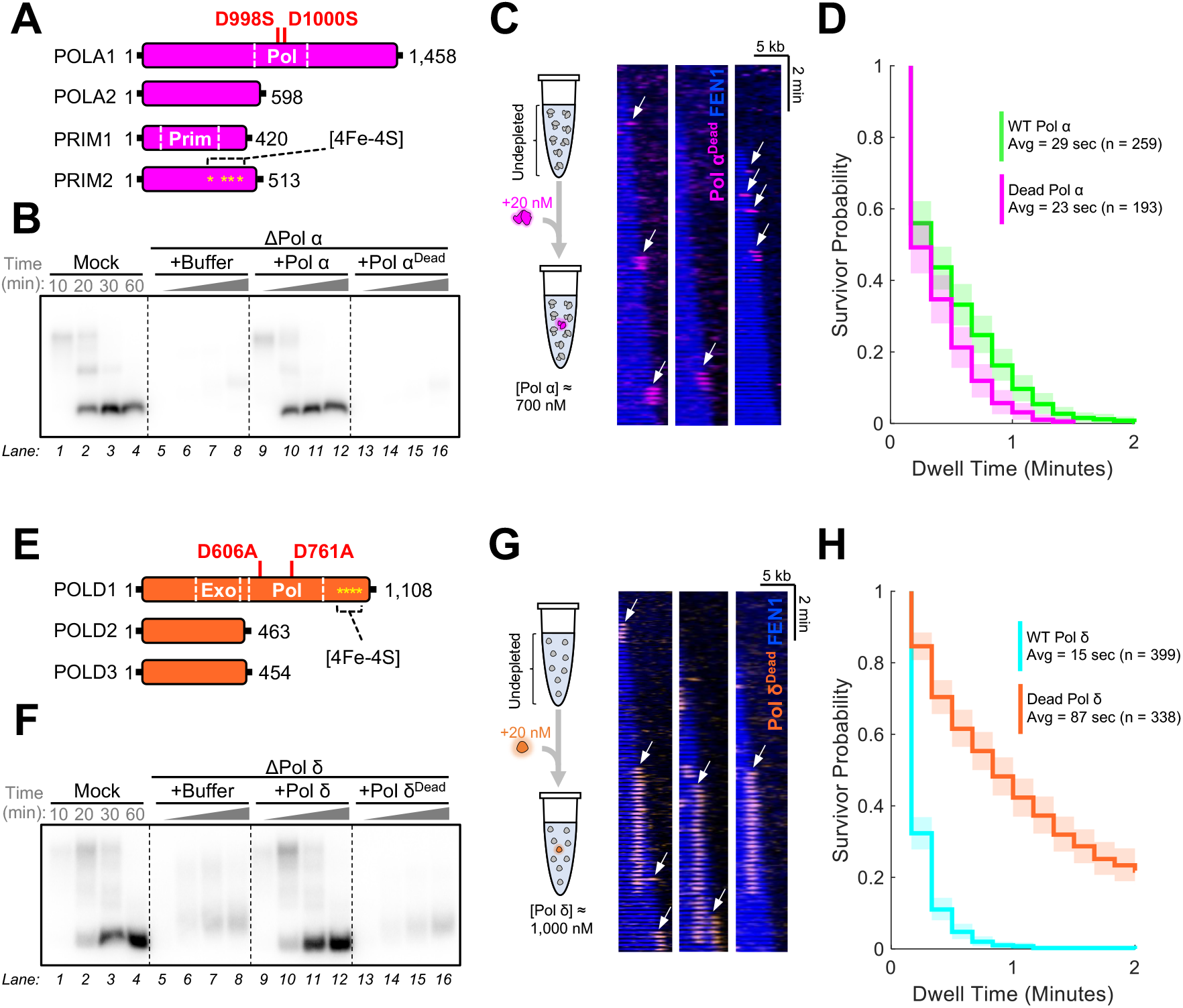
A Single Copy of Catalytically Dead Pol α/δ Does Not Impair Okazaki Fragment Synthesis. **(A)** Point mutations introduced into *Xenopus* Pol α to inactivate its DNA polymerase activity. **(B)** Ensemble plasmid replication assay where Pol α-depleted extract was supplemented with 100 nM of recombinant wildtype enzyme (Pol α) or catalytically inactive enzyme (Pol α^Dead^). Timepoints were collected at 10, 20, 30, and 60 minutes. **(C)** Representative kymograms from an experiment where 20 nM of fluorescent Pol α^Dead^ (magenta) was spiked in to undepleted extract containing ∼700 nM of endogenous Pol α. **(D)** Survivor probability curves of binding event dwell times for Pol α^Dead^ (magenta) versus wildtype Pol α (green). **(E)** Mutations introduced to inactivate *Xenopus* Pol δ. **(F)** Ensemble plasmid replication assay where Pol δ-depleted extract was supplemented with 100 nM of recombinant wildtype enzyme (Pol δ) or catalytically inactive enzyme (Pol δ^Dead^). **(G)** Representative kymograms from an experiment where 20 nM of fluorescent Pol δ^Dead^ (orange) was spiked in to undepleted extract containing ∼1,000 nM of endogenous Pol δ. **(H)** Survivor probability curves of binding event dwell times for Pol δ^Dead^ (orange) versus wildtype Pol α (cyan). In panels **(D)** and **(H)**, semi-transparent boxes depict the 95% Confidence Interval (CI) estimated via bootstrapping.

Similarly, we generated catalytically dead Pol δ (Pol δ^Dead^) by mutating two essential residues in the POLD1 subunit (D606A, D761A) as previously described^9^ (Figure 2E). We confirmed that Pol δ^Dead^ did not rescue replication in Pol δ-depleted extract (Figure 2F, S2B). Spiking 20 nM of fluorescently-labeled Pol δ^Dead^ into undepleted extract revealed that it also localizes to replication forks (Figure 2G). However, Pol δ^Dead^ resides on DNA much longer than the wildtype polymerase (Figure 2H), consistent with previous reports that catalytically dead human Pol δ is slow to dissociate from its template^9,27^. Strikingly, although Pol δ^Dead^ initially bound to the leading edge of the FEN1^mKikGR^ bubble, the fluorescently-labeled Pol δ^Dead^ remained stationary on DNA and did not travel with the advancing replication fork as marked by the growing edge of the FEN1^mKikGR^ tract (Figure 2G). This observation indicates that Pol δ is deposited behind the replication fork and corroborates the idea that Pol δ is not stably tethered to the replisome.

Taken together, the results of these spike-in experiments indicate that the replisome can tolerate the recruitment of a single copy of inactive Pol α or Pol δ without impairing lagging strand synthesis.

### Lagging Strand Synthesis Stops Shortly After Chasing with Catalytically Dead Pol α/δ

Previous studies proposed that the replisome may contain more than one copy of Pol α – one bound directly to CMG, and up to two copies tethered to the replisome via AND1 (Ctf4 in yeast)^5,17^. It is therefore possible, that in the Pol α^Dead^ spike-in experiment (Figure 2C), the recruitment of an inactive polymerase did not impair lagging strand synthesis because the replisome also contained a wildtype Pol α productively performing primer synthesis. If so, the trombone model predicts that the replisome should resist a chase with Pol α^Dead^ for several minutes, until all wildtype Pol α enzymes tethered to the replisome are replaced by Pol α^Dead^.

To test this prediction, we devised a single-molecule chase assay where we monitor lagging strand synthesis in real time by imaging FEN1^mKikGR^. We replicated DNA in undepleted extract for 10 minutes, then flushed the flow-cell with a “chase” extract and monitored the effect of the chase on lagging strand synthesis (Figure 3A). First, we chased with mock-depleted extract, which did not affect the growth rate of FEN1^mKikGR^ tracts (Figure 3B-C). Second, we chased with Pol α-depleted extract supplemented with 500 nM of Pol α^Dead^ (Figure 3D-F). Strikingly, chasing with Pol α^Dead^ caused FEN1^mKikGR^ tracts to stop growing ∼1 minute after the start of the chase (Figure 3E). Since ∼20 seconds are needed to completely flush our flow-cell, we conclude that all wildtype Pol α molecules at replication forks were replaced by Pol α^Dead^ within ∼40 seconds. Finally, we chased with Pol δ-depleted extract supplemented with 500 nM of Pol δ^Dead^ (Figure 3G-I). Similar to the Pol α^Dead^ chase, chasing with Pol δ^Dead^ abruptly stopped the growth of FEN1^mKikGR^ tracts within ∼1 minute (Figure 3H). Notably, we did not observe a subpopulation of forks that resisted the chase with Pol α^Dead^ or Pol δ^Dead^ (Figure 3F, 3I).

**Figure 3:**
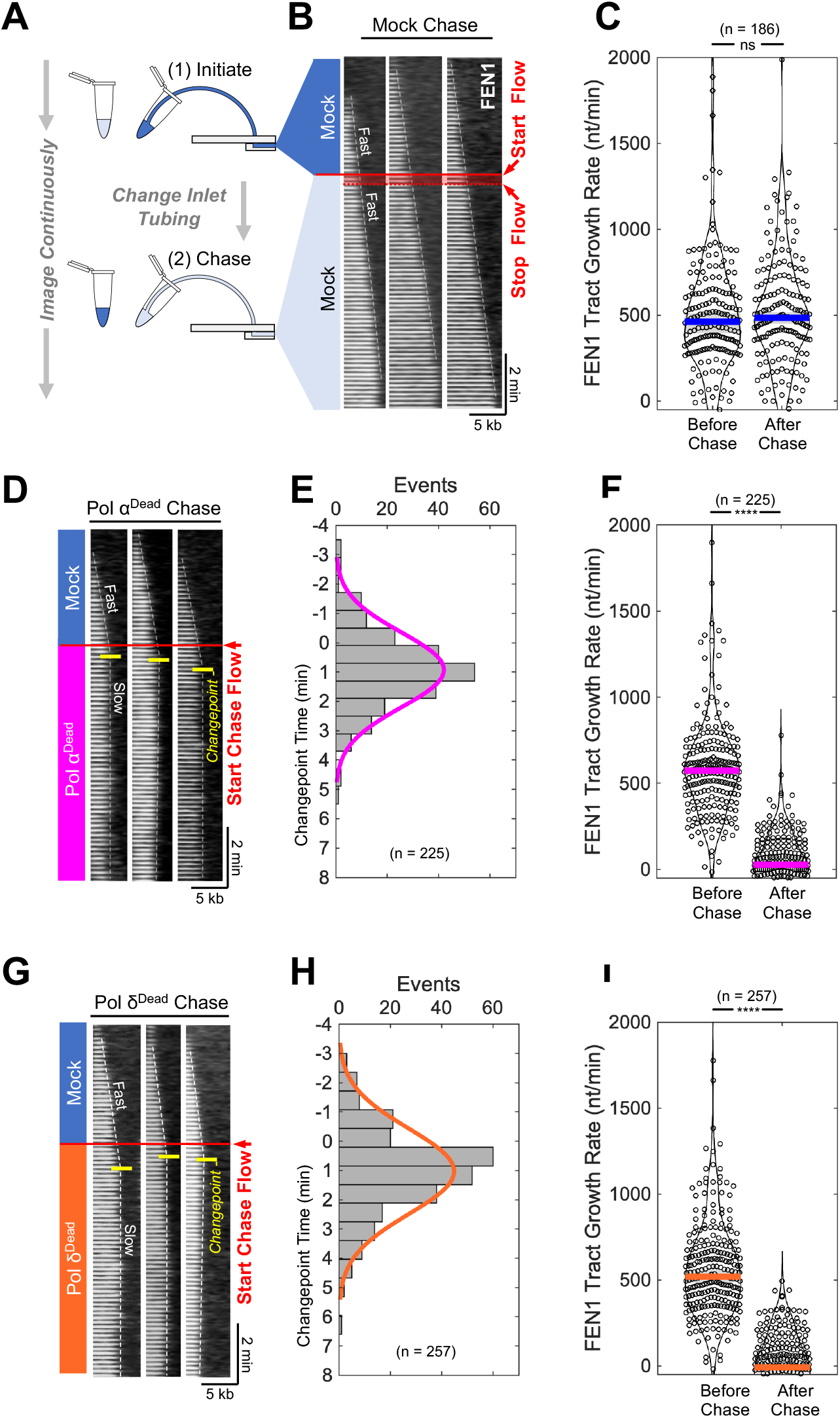
Chase with Catalytically Dead Pol α/δ Rapidly Impairs Okazaki Fragment Synthesis. **(A)** Diagram of the single-molecule chase experiment. **(B)** Representative kymograms of the lagging strand synthesis marker FEN1^mKikGR^ (gray) in the chase experiment with mock-depleted extract. The red line marks the start of the flow to pump the chase extract into the flow cell, which lasts ∼20 seconds. The dashed line marks the growing edge of the FEN1 tract. **(C)** The growth rate of the FEN1^mKikGR^ tract before and after the mock chase.**(D)** Representative kymograms of the lagging strand synthesis marker FEN1^mKikGR^ (gray) when the chase contained Pol α-depleted extract supplemented with 500 nM Pol α^Dead^. The dashed line marks the edge of the FEN1 tract fit by two straight lines with a changepoint (marked in yellow). **(E)** Time of the changepoint measured in the Pol α^Dead^ chase experiment. The changepoint was measured relative to the start of the chase flow as illustrated in panel **(D). (F)** The growth rate of the FEN1^mKikGR^ tract before and after the Pol α^Dead^ chase. **(G)** Representative kymograms of FEN1^mKikGR^ when the chase contained Pol δ-depleted extract supplemented with 500 nM Pol δ^Dead^. **(H)** Time of the changepoint measured in the Pol δ^Dead^ chase experiment. **(I)** The growth rate of the FEN1^mKikGR^ tract before and after the Pol δ^Dead^ chase. In panels **(C), (E), (F), (H)**, and **(I)** ‘n’ denotes the number of replication forks analyzed. In panels **(C), (F)**, and **(I)**, ‘ns’ denotes p>0.05, ‘****’ denotes p<0.0001. p-values were computed using the two-sample Kolmogorov-Smirnov test.

Taken together, these experiments unambiguously show that neither Pol α nor Pol δ are stably tethered to the replisome. Instead, our data indicate that new copies of Pol α and Pol δ must be frequently recruited to the replisome to support the synthesis of new Okazaki fragments.

### Leading and Lagging Strand Synthesis Are Not Intrinsically Coupled

It was previously proposed that by tethering the lagging strand polymerase to the replisome, the trombone model provides a way to coordinate leading and lagging strand synthesis^12,13^. Given our finding that Pol α and Pol δ are not stably tethered to the replisome, we asked whether leading and lagging strand synthesis are intrinsically coupled. To this end, we performed single-molecule chase experiments where we monitored (i) ssDNA accumulation by imaging Replication Protein A (RPA) tagged with mKikGR (RPA^mKikGR^) and (ii) CMG helicase dynamics by visualizing its subunit GINS labeled with Lumidyne555 (GINS^LD555^).

First, we chased with mock-depleted extract supplemented with DMSO (Figure 4A). During the mock chase CMG travelled fast (∼300 nt/min), comparable to CMG speed before the chase. There was very little RPA^mKikGR^ signal at the replication fork (Figure 4A), corresponding to normal leading/lagging strand synthesis where ssDNA does not accumulate behind CMG.

**Figure 4:**
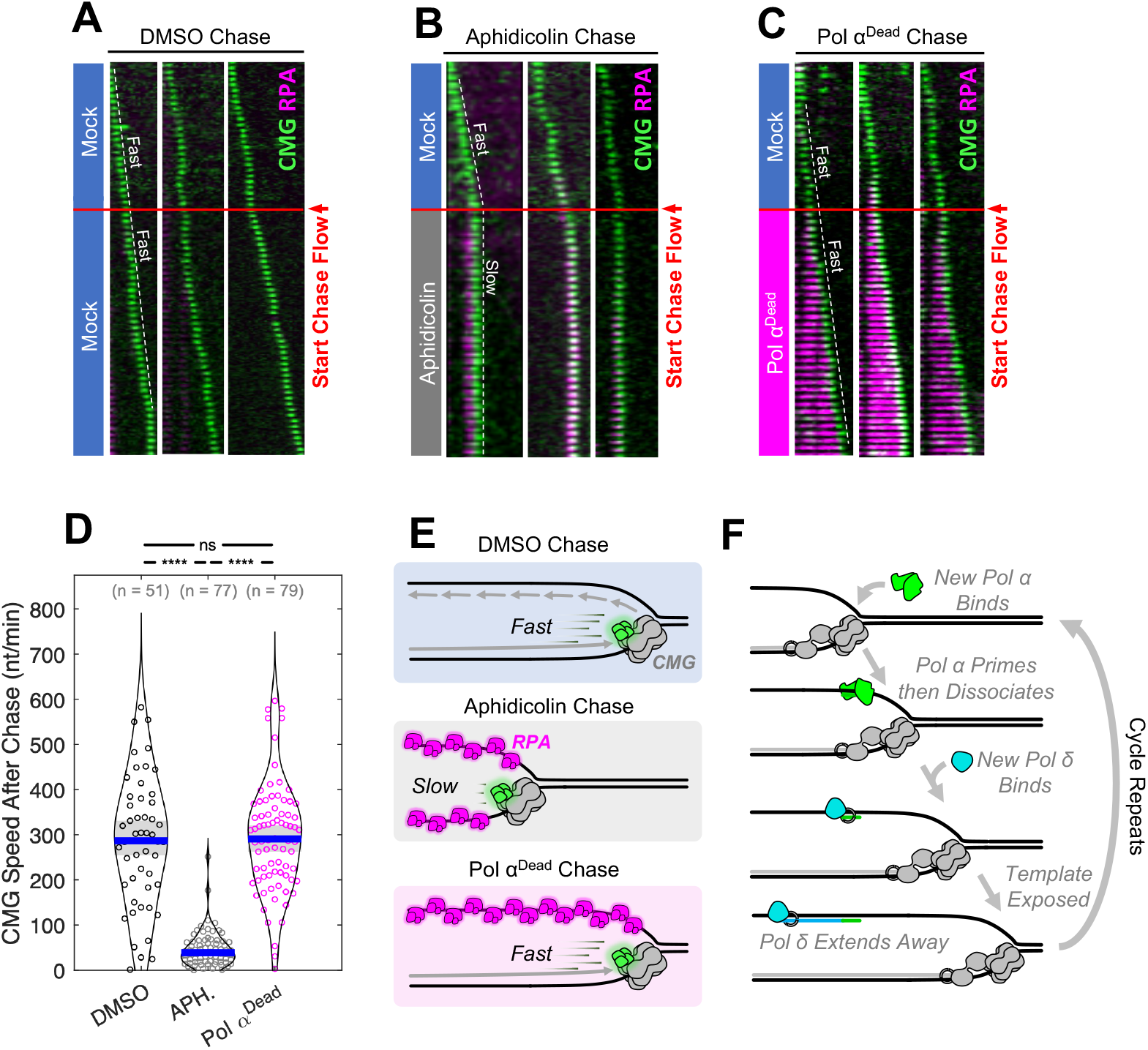
Leading and Lagging Strand Synthesis are Not Intrinsically Coupled. **(A)** Representative kymograms of CMG (green) and RPA^mKikGR^ (magenta) from a single-molecule chase experiment with mock-depleted extract. CMG was labeled by replacing endogenous GINS with fluorescently-labeled GINS^LD555^. The dashed line marks the progression of CMG. **(B)** Representative kymograms of CMG (green) and RPA^mKikGR^ (magenta) from an experiment where the chase extract was supplemented with 50 µM aphidicolin. **(C)** Representative kymograms of CMG (green) and RPA^mKikGR^ (magenta) from a chase with Pol α-depleted extract supplemented with 500 nM Pol α^Dead^. **(D)** The distribution of CMG speeds measured after the chase in experiments from panels (**A-C). (E)** Cartoons illustrating the replication defect after the chase in experiments from panels (**A-C). (F)** Illustration of the *distributive* mechanism of lagging strand synthesis.

Second, we chased with mock-depleted extract supplemented with 50 μM of aphidicolin, a B-family polymerase inhibitor that blocks both leading and lagging strand synthesis^28–30^ (Figure 4B). During the aphidicolin chase, the speed of CMG abruptly dropped to ∼50 nt/min and this was accompanied by the accumulation of RPA^mKikGR^ signal behind the helicase (Figure 4B). This corresponds to no DNA synthesis on the leading or lagging strand.

Finally, we chased with Pol α-depleted extract supplemented with 500 nM of Pol α^Dead^. During the Pol α^Dead^ chase we observed a long tract of RPA^mKikGR^ accumulating behind CMG (Figure 4C), confirming that Okazaki fragment synthesis was inhibited and ssDNA was accumulating on the lagging strand template. Notably, during the Pol α^Dead^ chase CMG travelled as fast as during the mock chase (Figure 4D), indicating that leading strand synthesis was not affected by blocking lagging strand synthesis (Figure 4E).

Taken together, these data indicate that the rate of leading strand synthesis is not intrinsically coupled to that of lagging strand synthesis.

## DISCUSSION

### Pol α and Pol δ Function *Distributively* During Lagging Strand Synthesis

The data presented here support a *distributive* mechanism of lagging strand synthesis in eukaryotes (Figure 4F). First, Pol α docks onto CMG as shown by a recent cryo-EM study^5^, enabling Pol α to engage the lagging strand template. We find that Pol α resides at the fork long enough for CMG to unwind 200-250 nt of template DNA (Figure 1D), indicating that new Pol α molecules must be recruited for priming new Okazaki fragments. Second, Pol δ is recruited to the newly synthesized primer. We find that Pol δ is deposited behind the fork (Figure 2G), suggesting that Pol δ synthesizes the lagging strand in the opposite direction of fork progression (Figure 4E). Similar to Pol α, this necessitates the recruitment of new Pol δ molecules to extend new Okazaki fragments. Finally, continued DNA unwinding by CMG exposes more of the lagging strand template, facilitating the recruitment of new molecules of Pol α and Pol δ for new rounds of *distributive* Okazaki fragment synthesis.

The *distributive* mechanism has several potential benefits. (i) It renders lagging strand replication robust to the occasional incorporation an inactive Pol α/δ into the replisome (Figure 2C, 2G). Notably, since both Pol α and Pol δ contain Fe-S clusters that are essential for their enzymatic function^31–33^, endogenous or exogenous oxidative stress may inactivate a fraction of all available polymerases^34^. (ii) It does not require active mechanisms to remove defective Pol α/δ from elongating replisomes. (iii) It does not strictly require the lagging strand template to form loops and obviates the need for any additional machinery to manage such loops. (iv) It could allow replisomes to more easily pass through cohesin rings. CMG alone has a diameter of 10-15 nm ^35^, so the presence of Pol α (12-15 nm in diameter)^7^ and Pol δ (7-10 nm in diameter)^36^ in addition to other replisome subunits may significantly impede the ability of the replisome to pass through a cohesin ring (30-40 nm in diameter)^37^.

What causes Pol α to dissociate from the replisome? In the absence of nucleotides, Pol α forms a stable complex with CMG on forked DNA, and this interaction is mediated by multiple CMG-Pol α contacts^5^. Notably, mutating just one of these interfaces destabilizes the CMG-Pol α interaction and reduces priming frequency *in vitro*^5^. Since Pol α undergoes major conformational changes during primer synthesis^7,8^, we speculate that these structural rearrangements disrupt Pol α’s interaction with CMG, triggering its dissociation from the replisome. Future studies capturing CMG-Pol α structures at different stages of primer synthesis are needed to test this prediction.

What causes Pol δ to dissociate from the replisome? Direct visualization of Pol δ^Dead^ reveals that Pol δ molecules are deposited on the lagging strand template behind advancing forks (Figure 2G). This suggests that Pol δ extends Okazaki fragments in the opposite direction of fork progression. Pol δ’s dissociation from template DNA could be triggered by its intrinsically low processivity^27^, or by its collision with the previous Okazaki fragment^11^. We propose that during lagging strand synthesis, Pol δ acts independently from the replisome, similar to how it functions during homologous recombination^38^ and break-induced replication^39^.

### Reconciling Our Observations with Previous Reports

A recent study used single-particle tracking to track Pol α/δ in live yeast and concluded that they bind chromatin long enough to support the synthesis of many consecutive Okazaki fragments^18^. However, it was not clear if the chromatin binding events measured in that study occurred at actively elongating replisomes. We find that Pol α resides at active forks for only ∼0.5 minutes compared to >2 minutes at stalled replisomes (Figure S4A). We surmise that some of the long-lived Pol α binding events reported *in vivo*^18^ occurred at stalled forks. In the case of Pol δ, it is unclear if the long-lived chromatin binding events measured *in vivo*^18^ reflect Pol δ’s binding at replication forks or Pol δ participating in DNA repair^38–40^. In human cells, Pol δ colocalizes with PCNA for only ∼4 seconds^9^ (corresponding to a fork travelling ∼130 nucleotides in human cells), consistent with the *distributive* mechanism of lagging strand synthesis.

Another study used a minimal *in vitro* reconstitution of the yeast replisome and found that single molecules of Pol α/δ travel with CMG long enough to synthesize >100 Okazaki fragments^17^. Notably, that study used polymerase concentrations 10-100x lower than in yeast/human cells^24,41^, which cause ATR checkpoint activation in live yeast^42,43^ and in nuclear extract (Figure S1J, S1M). We find that across physiologically relevant concentrations Pol α/δ function *distributively* (Figure 1D, 1F). Interestingly, the same study found that a polymerase-dead but primase-proficient Pol α mutant (equivalent to Pol α^Dead^ used here, Figure 2A) supported normal DNA replication *in vitro*^17^. This contrasts with our observation that Pol α^Dead^ cannot support replication in nuclear extract (Figure 2B). We surmise that RNA primers synthesized by Pol α^Dead^ are unstable in the nucleoplasm (and likely *in vivo*) because of RNases or other enzymes. Contrary to the previous report^17^, we conclude that Pol α’s DNA polymerase activity is essential for lagging strand synthesis.

Previous studies found that decreasing Pol α’s concentration *in vitro*^3^ or *in vivo*^42^ resulted in longer lagging strand fragments, which is difficult to reconcile with the idea that Pol α resides at the replisome for several Okazaki fragments. Instead, the inverse relationship between Okazaki fragment length and Pol α concentration is readily explained by the *distributive* mechanism supported by our data: low Pol α concentrations result in less efficient Pol α recruitment to forks and thus cause less frequent priming.

### Leading Strand Synthesis and Lagging Strand Synthesis Are Not Intrinsically Coupled

*In vitro*, fast leading strand synthesis and DNA unwinding can proceed in the absence of the lagging strand machinery^3,4^. We show that blocking lagging strand synthesis does not alter the rate of CMG progression in nuclear extract (Figure 4D). This contrasts with the “dead-man’s-switch” mechanism, where impairing leading strand synthesis immediately slows down the replicative DNA helicase^44^. Interestingly, the “dead-man’s switch” holds for prokaryotic^45,46^, and eukaryotic^47^ replicative helicases despite the fact that they evolved independently^48^. In conjunction with work from other labs, our findings indicate that the lack of intrinsic coupling between leading and lagging strand synthesis is an example of convergent evolution across the domains of life.

It was previously thought that leading and lagging strands were coordinated to prevent ssDNA accumulation on the lagging strand template. Our experiments reveal that there is no such coupling. Instead, our data support a model where the lagging strand machinery is rapidly recruited to forks ensuring that Okazaki fragment synthesis keeps up with the rate at which CMG unwinds DNA.

Although there is no intrinsic coupling between leading and lagging strand synthesis, recent studies suggest that external mechanisms may help coordinate these processes in cells. For example, treating human cells with the Pol α inhibitor CD437 leads to ATR activation and slows down replication forks, in part via fork reversal^49^. This suggests that ATR senses impaired lagging strand synthesis and restrains fork progression.

### Limitations

Immunodepleting Pol δ co-depleted ∼90% of Topoisomerase II α (Topo II α), causing a defect in plasmid decatenation (Figure S4B-C) that was fully rescued by 200 nM of recombinant *Xenopus* Topo II α (Figure S4D). To mitigate this, we supplemented Pol δ depleted extract with 200 nM recombinant Topo II α in all experiments. We considered the possibility that stretching the template DNA may disfavor the formation of trombone-like loops^12,13^. To this end, we measured the residence time of Pol α/δ on replicating singly-tethered DNA molecules which are not under tension (Figure 1B, dotted circle). We found that Pol α and Pol δ resided on single-tethered DNA for an average of 43 seconds (Figure S4E-G) and 18 seconds (Figure S4H-J) respectively – consistent with the *distributive* mechanism. The residence time measured on single-tethered DNA is slightly longer than that measured on double-tethered DNA (Figure S4F-G, S4I-J). We likely over-estimate Pol α/δ’s residence time on single-tethered DNA because distinct binding events sometimes overlap in a diffraction-limited spot.

The average residence time of Pol α at active forks is consistent with one Pol α molecule priming one Okazaki fragment. However, we cannot exclude the possibility that occasionally, one Pol α molecule may prime two consecutive Okazaki fragments, as we cannot directly measure individual lagging strand fragments in our single-molecule assay. Similarly, we cannot rule out a model in which more than one Pol δ molecule is needed to extend each Okazaki fragment, as proposed previously^9^.

## Acknowledgments

We thank Prof. Karlene Cimprich, Prof. Joanna Wysocka, Prof. Johannes Walter, Prof. David Pellman, Prof. Riki Terui, and Dr. Gongshi Bai for suggestions based on their critical reading of the manuscript. We thank Scott Berger and Jinho Park for valuable discussions during the preparation of this manuscript. We thank Prof. Johannes Walter for the generous gift of an initial batch of anti-POLA1 serum. We thank Prof. James Dewar for the generous gift of affinity-purified anti-TOP2A antibodies. We thank Prof. Tatsuro Takahashi for the generous gift of sera against subunits of the FACT complex. We thank Dr. Tomasz Swigut for assistance with bioinformatic analysis searching for the POLD4 subunit in *Xenopus*. L.D.L was supported in part by the NIH T32GM007276. G.C. is supported by an NSF CAREER Award (2144481), an NIGMS R35 award (GM147060), and an American Cancer Society seed grant (228425).

## Author Contributions

L.D.L. and G.C. conceived the project; L.D.L. carried out all experiments; G.C. and L.D.L. wrote MATLAB code to analyze single-molecule imaging movies; L.D.L. analyzed all data; L.D.L. and G.C. wrote the manuscript.

## SUPPLEMENTARY FIGURE CAPTIONS

**Figure S1:**
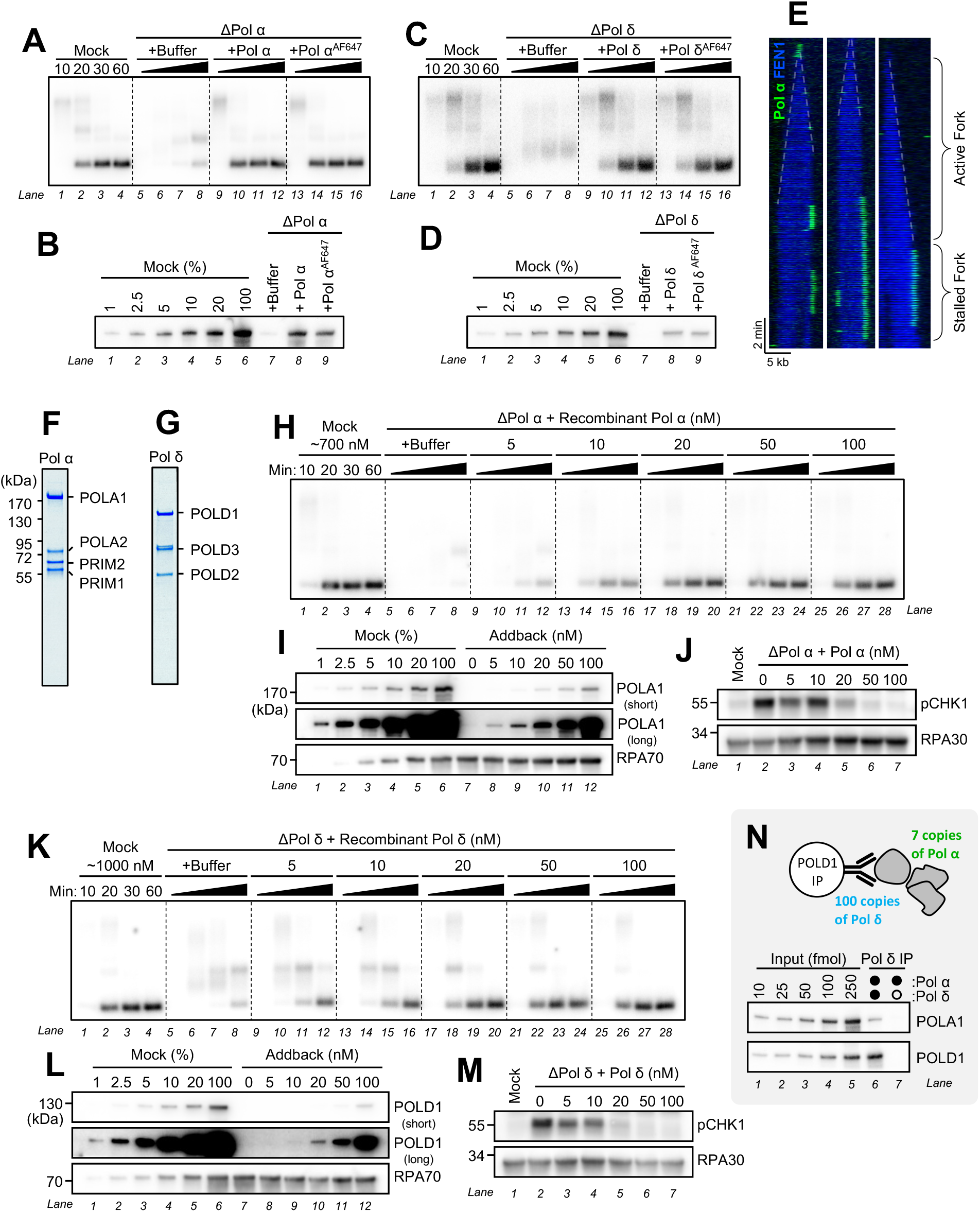
Validation of Recombinant Pol α/δ and Other Reagents. **(A)** Ensemble plasmid replication assay comparing the ability of Pol α and Pol α^AF647^ to support DNA replication in Pol α-depleted extract. **(B)** Immunoblot corresponding to the experiment in panel **(A). (C)** Ensemble plasmid replication assay comparing the ability of Pol δ and Pol δ^AF647^ to support DNA replication in Pol δ-depleted extract. **(D)** Immunoblot corresponding to the experiment in panel **(C). (E)** Representative kymograms Pol α^AF647^ binding at stalled replication forks. Dashes lines mark the growing edge of active forks. **(F)** Coomassie gel of purified recombinant Pol α used in this study. **(G)** Coomassie gel of purified recombinant Pol δ used in this study. **(H)** Replication assay depicting Pol α titration. **(I)** POLA1 immunoblot corresponding to the experiment in panel (**H). (J)** Phospho-CHK1 immunoblot for the experiment from panel (**H). (K)** Replication assay depicting Pol δ titration. **(L)** POLD1 immunoblot corresponding to the experiment in panel **(K). (M)** Phospho-CHK1 immunoblot for the experiment from panel (**K). (N)** Co-immunoprecipitation of Pol α with Pol δ.

**Figure S2:**
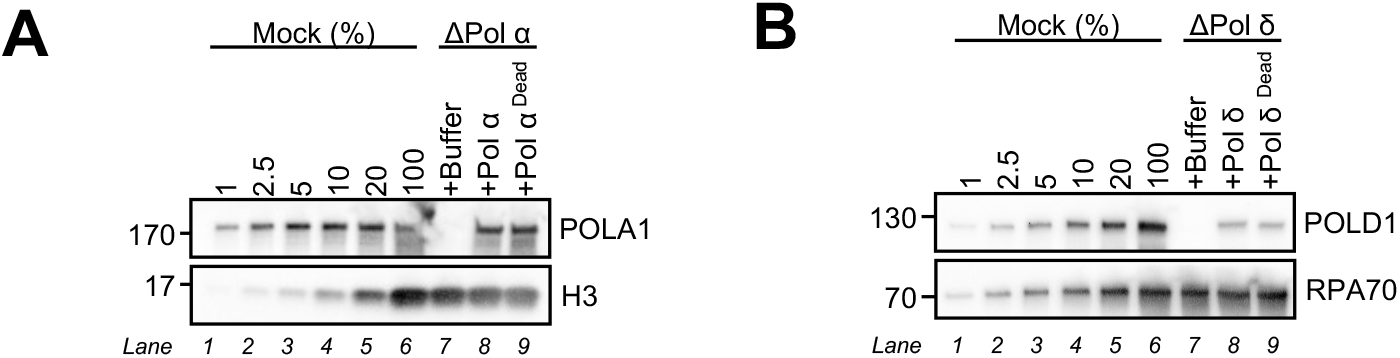
Validation of Catalytically Dead Pol α/δ Recombinant Proteins. **(A)** POLA1 immunoblot corresponding to the experiment from Figure 2B. **(B)** POLD1 immunoblot corresponding to the experiment from Figure 2F.

**Figure S3:**
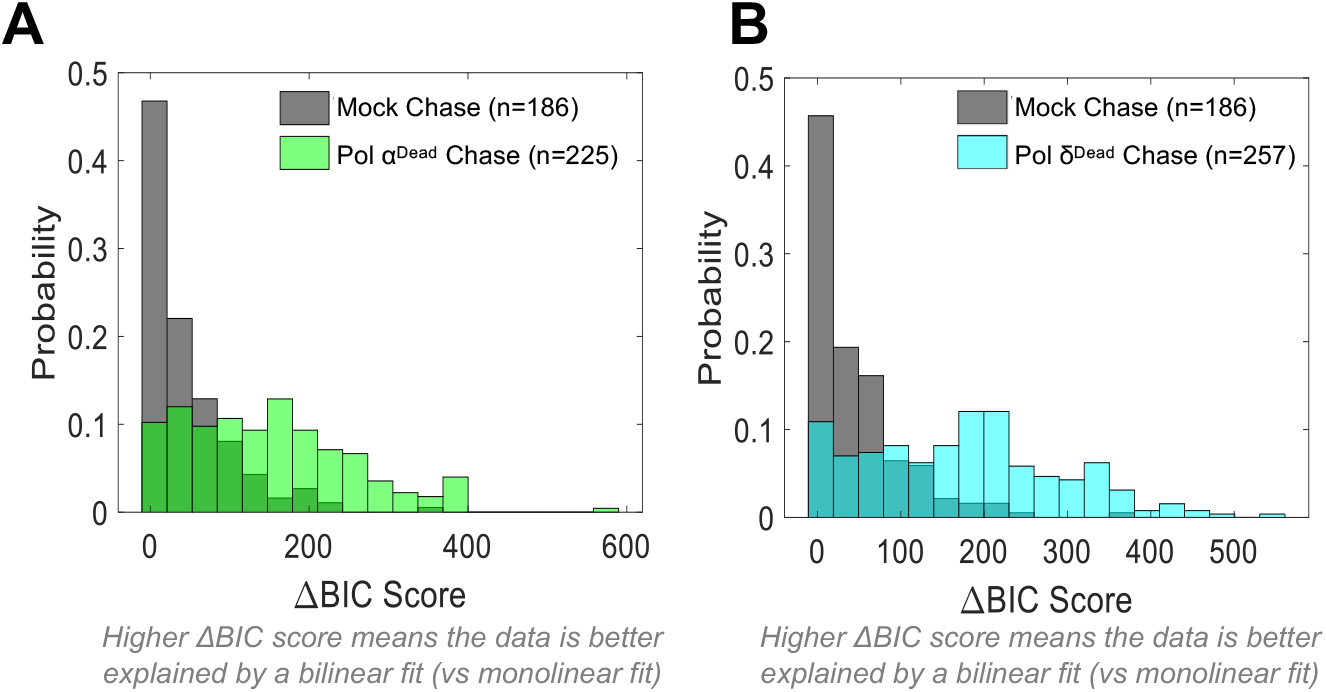
Validation of Changepoint Algorithm Used to Analyze Chase Experiments. **(A)** In the Pol α^Dead^ chase experiment, an automated thresholding algorithm was used to determine the position of the FEN1 tract edge, as shown in Figure 3D. The FEN1 edge position was fit to a monolinear fit or a bilinear fit. For each fit, a Bayesian Information Criterion (BIC) score was computed: BIC_monolinear_ and BIC_bilinear_. BIC quantifies how well the model (fit) explains the data and penalizes overfitting. A fit with a lower BIC score is considered to better explain the data. We computed ΔBIC = BIC_monolinear_ - BIC_bilinear_: higher ΔBIC values indicate the bilinear model is a better fit to the data. ΔBIC values are consistently higher for FEN1 tracts from the Pol α^Dead^ chase (green) compared to mock (gray), indicating that the growth of the FEN1 tract was better explained by a bilinear fit. **(B)** ΔBIC scores comparing FEN1 tracts from the Pol δ^Dead^ chase (cyan) versus mock (gray).

**Figure S4:**
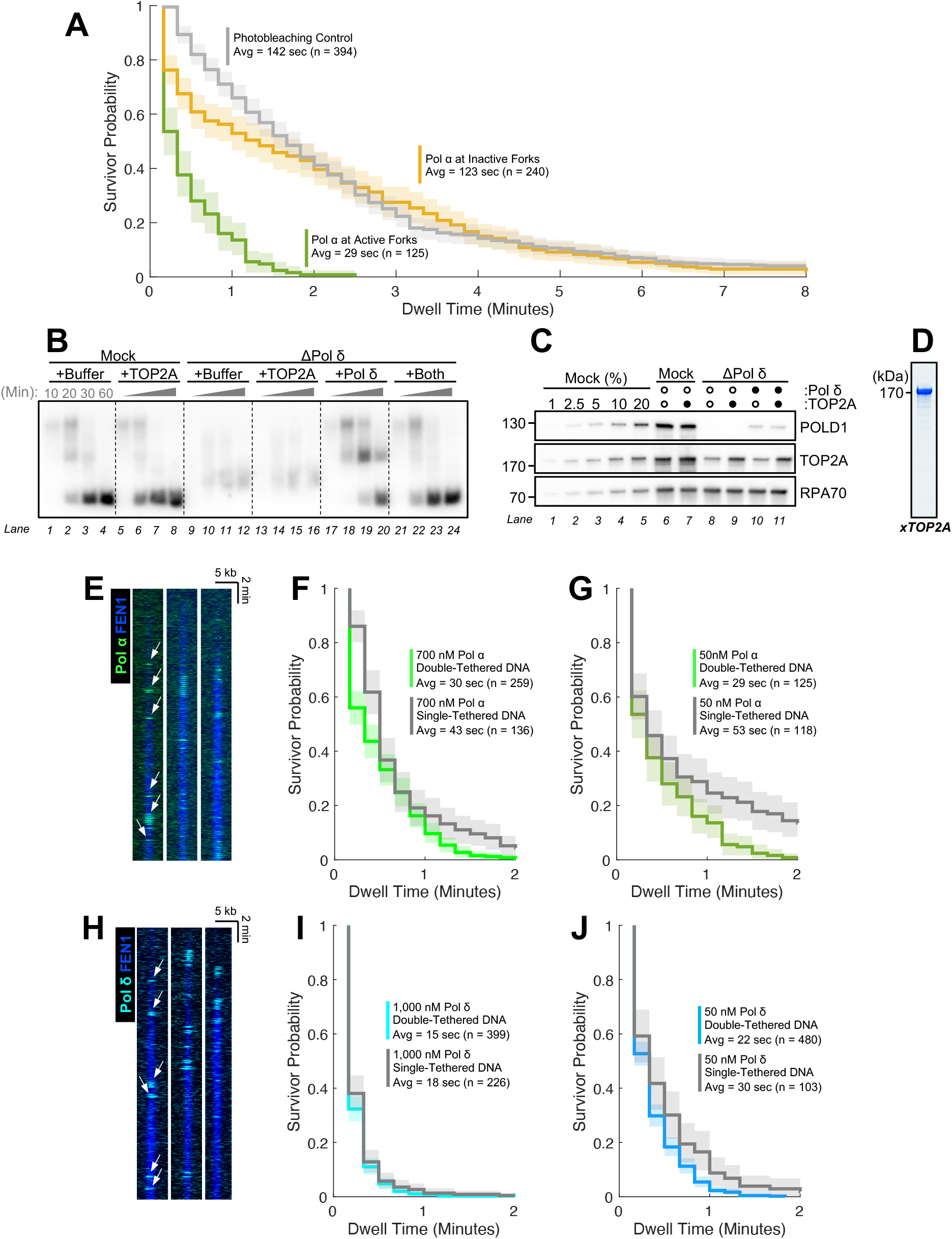
Additional Controls and Reagent Validation. **(A)** Survivor probability curves of dwell times for Pol α^AF647^ binding events at active forks (green) and stalled forks (orange). Note that the duration of Pol α^AF647^ binding events at stalled forks is comparable to the photobleaching control (gray), indicating that our ability to measure the dwell time of Pol α^AF647^ stalled forks is limited by photobleaching and that most likely, Pol α^AF647^ resides at stalled replication forks longer than the estimated 123 seconds. **(B)** Ensemble plasmid replication assay where Pol δ-depleted extract was supplemented with 200 nM recombinant TOP2A, 100 nM recombinant Pol δ, or both. Timepoints were collected at 10, 20, 30, and 60 minutes. **(C)** Immunoblot of POLD1 and TOP2A corresponding to the experiment from panel **(B)** reveals that TOP2A is partially co-immunodepleted with Pol δ. **(D)** Coomassie gel of purified recombinant TOP2A used in this study. **(E)** Representative kymograms of Pol α^AF647^ (green) from single-tethered DNA molecules which are condensed in a diffraction-limited spot. The accumulation of FEN1^mKikGR^ signal (blue) indicates that these DNA molecules are being replicated. **(F)** Survivor probability curves of dwell times for Pol α^AF647^ binding events on single-tethered DNA (gray) versus double-tethered DNA (green) from an experiment where 20 nM of Pol α^AF647^ was spiked into undepleted extract (∼700 nM of endogenous Pol α). **(G)** Same as in panel **(F)**, but for an experiment where 5 nM of Pol α^AF647^ was spiked into Pol α-depleted extract supplemented with 45 nM of recombinant Pol α (50 nM of total Pol α). **(H)** Representative kymograms of Pol δ^AF647^ (cyan) and FEN1^mKikGR^ (blue) from single-tethered DNA molecules. **(I)** Survivor probability curves of dwell times for Pol δ^AF647^ binding events on single-tethered DNA (gray) versus double-tethered DNA (cyan) from an experiment where 20 nM of Pol δ^AF647^ was spiked into undepleted extract (∼1000 nM of endogenous Pol δ). **(J)** Same as in panel **(I)**, but for an experiment where 5 nM of Pol δ^AF647^ was spiked into Pol δ-depleted extract supplemented with 45 nM of recombinant Pol δ (50 nM of total Pol δ).

