## Supplemental Information (methods) for "Eukaryotic Lagging Strand Synthesis is Distributive"

**Supplemental Materials**

**Key Resources Table**
See Excel Spreadsheet

**Experimental Model and Subject Details**
**Xenopus Laevis**
Eggs from adult, female *Xenopus laevis* frogs (NASCO Healthcare, Cat# LM00531) were used to prepare egg extracts and the testes of adult, male *Xenopus laevis* frogs (NASCO Healthcare, Cat# LM00715MX) were used to prepare sperm chromatin. All animals were healthy, not subjected to previous procedures, and no animal husbandry was performed. Frogs were housed at the Boswell Small Amphibians Research Facility at Stanford School of Medicine in compliance with Institute Animal Care and Use Committee (IACUC) regulations and all experiments involving frogs were approved by the Stanford University APLAC (Administrative Panel on Laboratory Animal Care).

**Insect Cell Lines**
Sf9 cells (Expression Systems, Cat# 94-001S) and Tni cells (Expression Systems, Cat# 94-002S) were cultured in ESF 921 insect cell culture medium (Expression Systems, Cat# 96-001-01) at 27°C for baculovirus production and protein overexpression.

**Bacterial Cell Lines Used**
DH10EMBacY (Geneva Biotech, Cat# DH10EMBacY), DH5α (New England Biolabs (NEB), Cat# C2987H), and Rosetta (DE3) pLysS (Novagen, Cat# 70956-4) bacteria were cultured in LB broth (Thermo Fisher Scientific, Cat# BP9723) at 37$^{o}$C for plasmid production and protein overexpression.

**Methods Details**
Preparation of DNA Substrates for Single-Molecule Studies
All single molecule experiments used doubly tethered λ phage DNA. The 48.5 kb DNA substrate as purchased from NEB (#N3011S) has 12-nt 5' ssDNA overhangs on both ends. We used the Klenow Fragment (3′ to 5′ exo-) polymerase (NEB, Cat# M0212S) to fill in the overhangs with biotin-dCTP (Thermo Fisher Scientific, Cat# 19518018) and biotin-dGTP (PerkinElmer, Cat# NEL541001EA). The product is separated with electrophoresis and electro-eluted from the gel before use in tethering. A step-by-step detailed protocol for preparing biotinylated λ DNA is available in (Berger and Chistol, 2024).

**Xenopus Egg Extract Preparation**
*Xenopus laevis* egg extracts were prepared following published protocols (Lebofsky et al., 2009; Sparks & Walter, 2019). Frogs were primed with 50 IU of human chorionic gonadotropin (CHORULON) (Merck, Cat# 133754) 5-7 days before extract preparation. To induce ovulation, 500 IU of CHORULON was injected 20-22 hr before extract preparation. High-speed supernatant (HSS) was prepared from 5-6 adult female frogs, and nucleoplasmic extract (NPE) was prepared from 15-20 frogs. Sperm chromatin was purified from adult male frog testes and used for NPE preparation.

**Ensemble Replication Assay**
To validate antibodies and recombinant proteins, ensemble DNA replication experiments were performed following a detailed protocol described in a recent methods article (Berger and Chistol, 2024). Since all proteins of interest (POIs) in this study are dispensable for Mcm2-7 double-hexamer loading, all licensing reactions were performed with HSS depleted of POI.

**Protein Co-Immunoprecipitation Assay**
To test if Pol α and Pol δ form a complex in solution, 100 nM of each polymerase was mixed together in a 20 µL reaction made up of ELB-Sucrose, 0.05% NP-40S, and 0.1 mg/mL BSA. The mixture was incubated for 1 hour at room-temperature with gentle rotation. 10 µg of affinity-purified anti-POLD1 antibody was incubated with 60

μg Dynabeads Protein A (Invitrogen, Cat# 10002D) for 1 hour at room-temperature with gentle rotation. Beads were washed twice with ELB-Sucrose + 0.05% NP-40S and then incubated with the polymerase mixture and incubated for 1 hour at room-temperature with gentle rotation. Beads were washed three times with ice-cold ELB-Sucrose + 0.05% NP-40S. Bound proteins were eluted by boiling beads in 15 μL 1x Laemmli buffer (50mM Tris-HCl pH 6.8, 2% SDS, 10% glycerol, 0.1% bromophenol blue, 2.5% β-mercaptoethanol).

**Western Blotting and Coomasie Staining**

All samples were boiled in 1x Laemmli buffer and run on 4-15% Mini-PROTEAN TGX Precast Protein Gels (Bio-Rad #4561086) using Tris-Glycine-SDS Running Buffer (Boston BioProducts #BP-150). EZ-Run Prestained Rec Protein Ladder (Fisher Scientific #BP36031) or Precision Plus Protein Dual Color Standard (Bio-Rad #1610374EDU) was run alongside samples to infer protein band sizes. For Coomasie staining, protein gels were incubated with Optiblot Blue InstantBlue Coomassie Protein Stain (Abcam #ab119211) for at least 1 hour at room temperature with gentle shaking. For immunoblotting, proteins were transferred from the gel onto a PVDF membrane (VWR #10061-492) in transfer buffer (Fisher Scientific #NC9297917). Membranes were blocked in 1x TBST (1x Tris Buffered Saline (TBS) (25 mM Tris pH 7.4, 150 mM NaCl), 0.1% Tween 20) containing 5% (w/v) BSA for 1 hour at room temperature with gentle shaking. Following a wash in 1xTBST, the membrane was incubated with primary antibodies diluted in 1x TBST containing 5% (w/v) BSA overnight at 4$^{o}$C with gentle shaking. Membranes were washed three times with 1x TBST and incubated with secondary antibody diluted in 1x TBST + 5% BSA for 1 hour at room temperature with gentle shaking. Membranes were washed three times with 1x TBST, developed using SuperSignal West Pico PLUS Chemiluminescent Substrate (Fisher Scientific, Cat# PI34577), and imaged using Azure c600 (Azure Biosystems).

**Cloning and Plasmids**

To prepare an antigen consisting of *Xenopus laevis* POLA1 amino acids 1 – 340, the corresponding DNA sequence was PCR amplified out of *X. laevis* egg cDNA. The resulting PCR product was assembled into pET28b(+) using the NEBuilder HiFi Mix (NEB, Cat# E2621S), yielding plasmid pJP006.

To prepare an antigen consisting of *X. laevis* POLD1 amino acids 1 – 89, the corresponding DNA sequence was PCR amplified out of *X. laevis* egg cDNA. The resulting PCR product was assembled into pET28b(+) using the NEBuilder HiFi Mix (NEB, Cat# E2621S), yielding plasmid pLDL048.

To prepare full-length *X. laevis* Pol α individual subunits were first cloned out of cDNA. The *pola1* gene was PCR amplified out of cDNA and assembled into BamHI/HindIII linearized pLib, yielding plasmid pLDL152. The *pola2* gene was PCR amplified out of cDNA and assembled into BamHI/HindIII linearized pLib, yielding plasmid pLDL153. The *prim1* gene was PCR amplified out of cDNA using primers and assembled into BamHI/HindIII linearized pLib, yielding plasmid pLDL154. The *prim2* gene was PCR amplified out of cDNA and assembled into BamHI/HindIII linearized pLib, yielding plasmid pLDL155. To tag the N-terminus of POLA1 with a TwinStrep-3aaLinker-HRV3C-S6-10aaLinker cassette, the *pola1* gene was PCR amplified out of pLDL152 and the resulting PCR product was assembled with a geneblock (synthesized by IDT) into BamHI/HindIII linearized pLib yielding plasmid pLDL156. All four subunits were then cloned into a single plasmid using the biGBac multigene construction strategy (Weissmann et al., 2016). *twinstrep-3aalinker-hrv3c-s6-10aalinker-pola1* was PCR amplified out of LDL156 using primers oLDL108/oLDL109, *pola2* was PCR amplified out of pLDL153 using primers oLDL110/oLDL111, *prim1* was PCR amplified out of pLDL154 using primers oLDL112/oLDL113, and *prim2* was PCR amplified out of pLDL155 using primers oLDL114/oLDL117. The four resulting PCR products were assembled into SwaI linearized pBiG1a, yielding the complete insect expression plasmid pLDL157.

To prepare a polymerase dead *X. laevis* Pol α, essential aspartic acids in the DNA polymerase active site of POLA1 were mutated to serine as previously described for the *Saccharomyces cerevisiae* ortholog (Lewis et al., 2020). D996 and D998 from *S. cerevisiae* Pol1 were mapped to D998 and D1000 in *X. laevis* POLA1.

To prepare full-length *X. laevis* Pol δ individual subunits were first cloned out of cDNA. The *pold1* gene was PCR amplified out of cDNA and assembled with a *3xflag-3aalinker-hrv3c-s6-10aalinker* containing geneblock (synthesized by IDT) into pACEBac1, yielding plasmid pLDL148. The *pold2* gene was PCR amplified out of cDNA and assembled into pIDC, yielding plasmid pLDL079. The *pold3* gene was PCR amplified out of cDNA using primers and assembled into pIDC, yielding plasmid pLDL080. All three subunits were then cloned into a single plasmid using the MultiBac multigene construction strategy (Sari et al., 2016). The *pold3* gene was extracted from pLDL080 and inserted into linearized pLDL079, yielding pLDL081. The pLDL081 and pLDL148 vectors were fused by Cre-recombination, yielding pLDL149. The 3xFLAG tag was replaced by a TwinStrep tag via Gibson assembly, yielding the complete insect expression plasmid pLDL158.

To prepare a polymerase dead *X. laevis* Pol δ, essential aspartic acids in the DNA polymerase active site of POLD1 were mutated to alanine as previously described for the *Homo sapiens* ortholog (Drosopoulos et al., 2020). D602 and D757 from *H. sapiens* POLD1 were mapped to D606 and D761 in *X. laevis* POLD1.

To prepare full- length *X. laevis* Topo II α the gene for *top2a* was cloned out of cDNA and inserted into a PCR linearized pLib backbone that already contained the DNA sequence for an upstream *twinstrep-HRV3C* cassette, yielding the insect expression plasmid pLDL189.

To prepare full-length *X. laevis* GINS with an S6 tag, a pGC124 containing *psf3-10aaLinker-sort-6xhis* and *sld5* was used as a starting point. A *10aalinker-s6-6xhis* cassette was added to the C-terminus of *psf3*, yielding the *psf3-10aaLinker-s6-6xhis* and *sld5* containing plasmid pLDL112. The pLDL112 (psf3-*10aaLinker-s6-6xhis/sld5)* and pGC033 (p*sf1*/*psf2*) vectors were fused by Cre-recombination, yielding the complete GINS expression plasmid pLDL113.

Oligos:

| Primer | Purpose | Sequence |
|---|---|---|
| oLDL108 | biGBac CasI_for | aacgctctatggtctaaagatttaaatcgacctactccggaatattaatagatc |
| oLDL109 | biGBac CasI_rev | aaacgtgcaatagtatccagtttatttaaatggttatgatagttattgctcagcg |
| oLDL110 | biGBac CasII_for | aaactggatactattgcacgtttaaatcgacctactccggaatattaatagatc |
| oLDL111 | biGBac CasII_rev | aaacatcaggcatcattaggtttatttaaatggttatgatagttattgctcagcg |
| oLDL112 | biGBac CasIII_for | aaacctaatgatgcctgatgtttaaatcgacctactccggaatattaatagatc |
| oLDL113 | biGBac CasIII_rev | aaactaagctatgtgaaccgtttatttaaatggttatgatagttattgctcagcg |
| oLDL114 | biGBac CasIV_for | aaacggttcacatagcttagtttaaatcgacctactccggaatattaatagatc |
| oLDL117 | biGBac_Casω_rev | aaccccgattgagatatagatttatttaaatggttatgatagttattgctcagcg |

Plasmids:

| Plasmid | Description |
|---|---|
| pJP006 | Pol α antigen (POLA1; Amino acids 1 – 340-6xHis) |
| pLDL048 | Pol δ antigen (POLD1; Amino Acids 1 – 89 -6xHis) |
| pLDL113 | GINS-S6-6xHis |
| pLDL157 | TwinStrep-HRV3C-S6-Pol α |
| pLDL158 | TwinStrep-HRV3C-S6-Pol δ |
| pLDL171 | TwinStrep-HRV3C-S6-Pol $\alpha^{Dead}$ (POLA1; D998S and D1000S) |
| pLDL184 | TwinStrep-HRV3C-S6-Pol $\delta^{Dead}$ (POLD1; D606A and D761A) |
| pLDL189 | Twinstrep-HRV3C-Topoisomerase II α |

**Antibody Preparation**

Custom polyclonal antibodies were generated as previously described (Berger and Chistol, 2024). Briefly, the His6 tagged antigen sequence was cloned into pET28b(+) (see Cloning and Plasmids). The antigen was expressed in Rosetta (DE3) pLysS (Novagen, Cat# 70956-4) upon induction with 0.5-1.0 mM IPTG for 3 hrs at 37°C. Soluble antigens were purified in native buffer, whereas insoluble antigens were purified in denaturing buffer (see table and 'Recombinant Protein Expression and Purification' below). The Ni-NTA eluate (still containing 400 mM imidazole) was used to immunize rabbits. All antibodies used in this study were raised by Cocalico Biologicals.

All antibodies used in this study were affinity purified. 1 – 5 mg of purified antigen was cross-linked to 1 mL of AminoLink Plus Coupling Resin (Fisher Scientific, Cat# PI20501) according to the manufacturer's protocol. 10 – 30 mL of serum was incubated with the affinity resin for 30 minutes at room temperature. The resin was washed twice with 10 mL of 1x PBS + 500 mM NaCl, and twice again with 10 mL of 1x PBS. Next, antibodies bound to the affinity resin were eluted 5-6 times with 1.0 mL of 200 mM glycine-HCl pH2.5. Each elution was collected into an tube containing 0.1 mL of 1M Tris-HCl pH9.0, which immediately neutralized the glycine. Eluted fractions were pooled and dialyzed against 1 L of 1x TBS (50 mM Tris-HCl pH8.0, 150 mM NaCl) + 10% (w/v) sucrose. The IgG concentration was measured using a Nanodrop, normalized to 1 mg/mL via dilution or spin-concentration, aliquots were flash frozen in liquid nitrogen, and stored at -80°C.

| Target Protein (Rabbit #) | Antigen | Plasmid | Antigen Purification | Depletion Condition µg IgG per µL extract | Primary Ab. Conc. for WB |
|---|---|---|---|---|---|
| Anti-xGINS (SU289) | Whole GINS complex Psf2-10aa-sort-His6 | pGC127 (insect) | Native | 1:1 | 0.2 µg/mL |
| Anti-xPOLA1 (SU375) | (1-340 aa)-His6 | pJP006 (bacteria) | Native | 2:1 | 0.2 µg/mL |
| Anti-xPOLD1 (SU329) | (1-89 aa)-His6 | pLDL048 (bacteria) | Native | 1:1 | 0.2 µg/mL |

**Recombinant Protein Expression and Purification**

$RPA^{mKikGR}$, $FEN1^{mKikGR}$, and SFP were purified following previously published protocols (Loveland et al., 2012; Modesti, 2011; Yin et al., 2006).

Pol α was purified as follows. A bacmid was generated by transforming DH10EMBacY cells with [pLDL157] pBiG_TwinStrep-HRV3C-S6-Pol α, and the resulting bacmid was purified with the ZR BAC DNA Miniprep Kit (Zymo Research, Cat# D4049). Baculovirus was amplified in three 72-hour stages (P1, P2, P3) in Sf9 cells. TwinStrep-HRV3C-S6-Pol α was expressed in 1 L of Tni suspension culture by infection with 2 mL of P3 baculovirus for 48 hrs. Cells were harvested, pelleted by centrifugation, and flash frozen in liquid nitrogen. Cell pellets were thawed and resuspended in 100 mL of buffer N (25 mM HEPES-NaOH pH 7.5, 300 mM NaCl, 10% glycerol, 1 mM DTT – bubbled with argon gas to remove dissolved oxygen) supplemented with 1x Pierce Protease Inhibitor EDTA-free (ThermoFisher Scientific, Cat# A32965) and lysed by Dounce homogenization. Lysate was cleared by ultracentrifugation at 35,000 rpm for 45 min at 4°C in a Beckman Type 70 Ti rotor (Beckman Coulter). Cleared lysate was incubated with 1 mL of Strep-Tactin®XT 4®Flow resin (IBA Lifesciences, Cat# 2-5010-010) for 1 hr at 4°C. The resin was washed with 3x10 mL buffer N. The protein was eluted from the resin using 4 mL buffer N supplemented with 50 µg/mL HRV3C Protease for 2 hours at 4°C on a rotator, and flash frozen in liquid nitrogen. Eluent was thawed and, diluted with 2 mL of buffer A (25 mM HEPES-NaOH pH 7.5, 10% glycerol, 1 mM DTT) to reduce the NaCl concentration to 200 mM, and loaded on a Mono Q 5/50 GL column (Cytiva, Cat# 17516601). The protein was eluted in 20 CVs off the column with a linear gradient of 200 mM – 1000 mM NaCl in buffer A and eluted around ~400 mM NaCl. Peak fractions were pooled, spin concentrated to ~0.5 mL using the Amicon Ultra-4-10K column (Merck Millipore, Cat# UFC801024) applied to a Superdex 200 Increase 10/300 gel-filtration column (Cytiva, Cat# 28-9909-44) equilibrated in 25 mM HEPES-NaOH pH 7.5, 10% glycerol, 1 mM DTT and 150 mM NaCl. Peak fractions were pooled, spin concentrated to ~6 µM using the Amicon Ultra-4-10K column (Merck Millipore, Cat# UFC801024) frozen in liquid nitrogen and stored at −80 °C. Approximately 100 µg total protein was obtained from 1 L of Tni cells. The Pol $\alpha^{Dead}$ mutant was purified as described for the wild-type protein.

Pol δ was purified as follows. A bacmid was generated by transforming DH10EMBacY cells with [pLDL158] pBiG_TwinStrep-HRV3C-S6-Pol δ, and the resulting bacmid was purified with the ZR BAC DNA Miniprep Kit (Zymo Research, Cat# D4049). Baculovirus was amplified in three 72-hour stages (P1, P2, P3) in Sf9 cells. We initially attempted to express Pol δ in Tni Cells, however the POLD3 subunit was not recovered (presumably due to proteolysis). Instead, TwinStrep-HRV3C-S6-Pol δ was expressed in 2 L of Sf9 suspension culture by infection with 4 mL of P3 baculovirus for 65 hrs. Cells were harvested, pelleted by centrifugation, and flash frozen in liquid nitrogen. Cell pellets were thawed and resuspended in 200 mL of buffer N (25 mM HEPES-NaOH pH 7.5, 300 mM NaCl, 10% glycerol, 1 mM DTT – bubbled with argon gas to remove dissolved oxygen) supplemented with 1x Pierce Protease Inhibitor EDTA-free (ThermoFisher Scientific, Cat# A32965) and lysed by Dounce homogenization. Lysate was cleared by ultracentrifugation at 35,000 rpm for 45 min at 4°C in a Beckman Type

70 Ti rotor (Beckman Coulter). Cleared lysate was incubated with 1 mL of Strep-Tactin®XT 4®Flow resin (IBA Lifesciences, Cat# 2-5010-010) for 1 hr at 4°C. The resin was washed with 3x10mL buffer. The protein was eluted from the resin using 4 mL buffer N supplemented with 50 µg/mL HRV3C Protease for 2 hours at 4°C on a rotator, and flash frozen in liquid nitrogen. Eluent was thawed and, diluted with 2 mL of buffer A (25 mM HEPES-NaOH pH 7.5, 10% glycerol, 1 mM DTT) to reduce the NaCl concentration to 200 mM, and loaded on a Mono Q 5/50 GL column (Cytiva, Cat# 17516601). The protein was eluted in 20 CVs off the column with a linear gradient of 200 mM – 1000 mM NaCl in buffer A and eluted around ~350 mM NaCl. Peak fractions were pooled, spin concentrated to ~0.5 mL using the Amicon Ultra-4-10K column (Merck Millipore, Cat# UFC801024) applied to a Superdex 200 Increase 10/300 gel-filtration column (Cytiva, Cat# 28-9909-44) equilibrated in 25 mM HEPES-NaOH pH 7.5, 10% glycerol, 1 mM DTT and 150 mM NaCl. Peak fractions were pooled, spin concentrated to ~4 µM using the Amicon Ultra-4-10K column (Merck Millipore, Cat# UFC801024) frozen in liquid nitrogen and stored at −80 °C. Approximately 50 µg total protein was obtained from 2 L of Sf9 cells. The Pol $\delta^{Dead}$ mutant was purified as described for the wild-type protein.

Topo II α was purified as follows. A bacmid was generated by transforming DH10EMBacY cells with [pLDL189] pLib_TwinStrep-HRV3C-Topo II α, and the resulting bacmid was purified with the ZR BAC DNA Miniprep Kit (Zymo Research, Cat# D4049). Baculovirus was amplified in three 72-hour stages (P1, P2, P3) in Sf9 cells. TwinStrep-HRV3C-Topo II α was expressed in 1 L of Sf9 suspension culture by infection with 2 mL of P3 baculovirus for 65 hrs. Cells were harvested, pelleted by centrifugation, and flash frozen in liquid nitrogen. Cell pellets were thawed and resuspended in 100 mL of buffer N (25 mM HEPES-NaOH pH 7.5, 300 mM NaCl, 10% glycerol, 1 mM DTT ) supplemented with 1x Pierce Protease Inhibitor EDTA-free (ThermoFisher Scientific, Cat# A32965) and lysed by Dounce homogenization. Lysate was cleared by ultracentrifugation at 35,000 rpm for 45 min at 4°C in a Beckman Type 70 Ti rotor (Beckman Coulter). Cleared lysate was incubated with 1 mL of Strep-Tactin®XT 4®Flow resin (IBA Lifesciences, Cat# 2-5010-010) for 1 hr at 4°C. The resin was washed with 3x10mL buffer. The protein was eluted from the resin using 4 mL buffer N supplemented with 50 µg/mL HRV3C Protease for 2 hours at 4°C on a rotator, and flash frozen in liquid nitrogen. 2 mL of eluent was thawed and, diluted with 4 mL of buffer A (25 mM HEPES-NaOH pH 7.5, 10% glycerol, 1 mM DTT) to reduce the NaCl concentration to 100 mM, and loaded on a Mono S 5/50 GL column (Cytiva, Cat# 17516801). The protein was eluted in 20 CVs off the column with a linear gradient of 100 mM – 1000 mM NaCl in buffer A and eluted around ~300 mM NaCl. Peak fractions were pooled, spin concentrated to ~0.5 mL using the Amicon Ultra-4-10K column (Merck Millipore, Cat# UFC801024) applied to a Superdex 200 Increase 10/300 gel-filtration column (Cytiva, Cat# 28-9909-44) equilibrated in 25 mM HEPES-NaOH pH 7.5, 10% glycerol, 1 mM DTT and 150 mM NaCl. Peak fractions were pooled, spin concentrated to ~4 µM using the Amicon Ultra-4-10K column (Merck Millipore, Cat# UFC801024) frozen in liquid nitrogen and stored at −80 °C. Approximately 300 µg total protein was obtained from 1 L of Sf9 cells.

GINS was purified as follows. A bacmid was generated by transforming DH10EMBacY cells with [pLDL113] pACEBac1_GINS-S6-6xHis, and the resulting bacmid was purified with the ZR BAC DNA Miniprep Kit (Zymo Research, Cat# D4049). Baculovirus was amplified in three 72-hour stages (P1, P2, P3) in Sf9 cells. GINS-S6-6xHis was expressed in 1/2 L of Tni suspension culture by infection with 2 mL of P3 baculovirus for 48 hrs. Cells were harvested, pelleted by centrifugation, and flash frozen in liquid nitrogen. Cell pellets were thawed and resuspended in 100 mL of buffer N (25 mM HEPES-NaOH pH 7.5, 500 mM NaCl, 10% glycerol, 1 mM DTT, 1 mM PMSF, 20 mM Imidazole) supplemented with 1x Pierce Protease Inhibitor EDTA-free (ThermoFisher Scientific, Cat# A32965) and lysed by Dounce homogenization. Lysate was cleared by ultracentrifugation at 35,000 rpm for 45 min at 4°C in a Beckman Type 70 Ti rotor (Beckman Coulter). Cleared lysate was incubated with 1 mL of Strep-Tactin®XT 4®Flow resin (IBA Lifesciences, Cat# 2-5010-010) for 1 hr at 4°C. The resin was washed with 3x10 mL buffer N. The protein was eluted from the resin using 5x0.5mL buffer N supplemented with 300 mM Imidazole, and flash frozen in liquid nitrogen. Eluent was thawed and, diluted with buffer A (25 mM HEPES-NaOH pH 7.5, 10% glycerol, 1 mM DTT) to reduce the NaCl concentration to 150 mM, and loaded on a Mono Q 5/50 GL column (Cytiva, Cat# 17516601). The protein was eluted in 20 CVs off the column with a linear gradient of 150 mM – 800 mM NaCl in buffer A and eluted around ~300 mM NaCl. Peak fractions were pooled, spin concentrated to ~0.5 mL using the Amicon Ultra-4-10K column (Merck Millipore, Cat# UFC801024) applied to a Superdex 200 Increase 10/300 gel-filtration column (Cytiva, Cat# 28-9909-44) equilibrated in 25 mM HEPES-NaOH pH 7.5, 10% glycerol, 1 mM DTT and 150 mM NaCl. Peak fractions were pooled, spin concentrated to ~3 μM using the Amicon Ultra-4-10K column (Merck Millipore, Cat# UFC801024) frozen in liquid nitrogen and stored at −80 °C. Approximately 2 mg total protein was obtained from 1/2 L of Tni cells.

**Fluorescent Labeling Using SFP Synthase**

For AF647 labeling, 1 mg of Alexa Fluor 647 C2 maleimide (Thermo Fisher Scientific, Cat # A20347) was reacted with 2 mg CoA-SH (Sigma-Aldrich, Cat # 234101-100MG) in 100 mM sodium phosphate pH 7.0 for 1 hour in the dark on a rotator at room temperature. The resulting product was separated from unreacted dye and unreacted CoA-SH by preparative HPLC using a reverse-phase Eclipse XDB-C18 column (PN 977250-102, 21.2 x 250 mm cartridge, 7 μm). The sample was run with a gradient of 0–50% (vol/vol) acetonitrile in 0.1% (vol/vol) TFA/water over 30 min at 10 mL/min. The AF647-CoA product eluted around ~36% acetonitrile. Peak fractions were pooled, lyophilized, and resuspended to a final concentration of 1 mM in DMSO. For LD555 and LD655 labeling, LD555-CoA and LD655-CoA were purchased from Lumidyne Technologies.

Site-specific fluorescent labeling of the S6 tag was performed in between the chromatography steps described above. Immediately following ion-exchange chromatography the molar concentration of the POI was estimated by the integrated intensity of the Abs280 reading on the FPLC chromatogram using a theoretical extinction coefficient. The POI was spin concentrated to ~0.5 mL and supplemented with 10 mM $MgCl_2$, a 2-fold molar excess of SFP synthase and a 10-fold molar excess of CoA conjugated dye. Reactions were incubated for 1 hour in the dark on a rotator at room temperature, then applied to a Superdex 200 Increase 10/300 gel-filtration column to separate unreacted fluorophore and SFP from the POI. The peak fractions were pooled and concentrated. Protein labeling efficiency was estimated based on absorbance measurements at 280 nm and the absorption peak for the respective dye. The biochemical activity of the labeled protein was tested following a published detailed protocol (Berger and Chistol, 2024).

**Single Molecule Replication Assay**

All single molecule experiments were conducted following a detailed protocol (Berger and Chistol, 2024), except for the following modifications. In the λ DNA tethering step, 20 pM of streptavidin$^{AF647}$ (Thermo Fisher Scientific, Cat# S21374) was added to the streptavidin solution. Streptavidin$^{AF647}$ spots were used as a fiducial marker for focusing immediately before starting imaging. After λ DNA tethering, DNA was licensed in HSS for 10 minutes. In this study, Replication Mix (10 μL of NPE, 10 μL of HSS, 3 - 6 μL of ELB-Sucrose, 1 μL of 600 ng/μL pBlueScript carrier plasmid DNA, 1 μL of ATP regeneration Mix, 1 μL of 60 μM FEN1$^{mKikGR}$ or 1 μL of 11 μM RPA$^{mKikGR}$, and 1- 4 μL of 150-900 nM fluorescently labeled protein of interest = 30 μL total) was flowed into the microfluidic chamber at a rate of 10 μL/min. The reaction was then imaged continuously (10-30 sec/frame) without washing out the free fluorescently labeled protein. Thus, fluorescent protein (5-20 nM) was present in the replication reaction during the entire duration of the experiment. Optimal replication efficiency was achieved when the single-molecule replication reaction contained equal volumes of NPE, HSS, and buffer (Yardimci et al., 2012). The concentrations of proteins examined in this study were estimated using quantitative western blots (see table below).

| *Protein of Interest* | **Concentration in NPE (nM)** | **Concentration in Replication Reaction (nM)** |
|---|---|---|
| **Pol α** | ~2,100 nM | ~700 nM |
| **Pol δ** | ~3,000 nM | ~1,000 nM |

For single molecule “chase” experiments the same procedure described above was repeated through the licensing step. Following licensing Replication Mix was flowed into the microfluidic chamber to initiate the reaction. After the flow completed, the inlet tubing was changed to a Chase Mix without starting flow. The Streptavidin$^{AF647}$ spots were used as a fiducial marker for focusing immediately before starting imaging. 20 – 30 minutes into the imaging acquisition the Chase Mix was flowed into the microfluidic chamber during continuous imaging.

For single-color imaging, FEN1$^{mKikGR}$ (488 nm laser excitation) was used. The following settings were used for imaging FEN1$^{mKikGR}$: TIRF arm set to 9000 (~67°), 500 ms exposure, 10 s/frame, 8% 488 nm laser power

(~60 µW power out of the objective, ~0.6 W/cm$^2$ power density), 100 EM Gain. Given these exposure durations, 12 fields of view (FOVs) could be sequentially imaged during the 10 s/frame interval.

For two-color imaging, FEN1$^{mKikGR}$ (488 nm laser excitation) and POI labeled with Alexa Fluor 647 (640 nm excitation) were used. The following settings were used for imaging Alexa Fluor 647: TIRF arm set to 9000 (~67°), 500 ms exposure, 10 s/frame, 25% 640 nm laser power (~380 µW power out of the objective, ~3.8 W/cm$^2$ power density), 100 EM Gain. FEN1$^{mKikGR}$ was imaged using the following settings: TIRF arm set to 9000 (~67°), 500 ms exposure, 10 s/frame, 8% 488 nm laser power (~60 µW power out of the objective, ~0.6 W/cm$^2$ power density), 100 EM Gain. RPA$^{mKikGR}$ was imaged using the following settings: TIRF arm set to 9000 (~67°), 500 ms exposure, 10 s/frame, 10% 488 nm laser power (~68 µW power out of the objective, ~0.68 W/cm$^2$ power density), 100 EM Gain. Given these exposure durations, 6 fields of view (FOVs) could be sequentially imaged during the 10 s/frame interval.

**Single-Molecule Data Analysis**

Each replication initiation experiment imaged 6-12 fields of view (FOVs) over the course of 45-60 minutes with a resolution of 10-30 seconds/frame. Each FOV typically contained 50-200 DNA molecules stretched to ~80% of their contour length. A suite of MATLAB scripts written in house were used to select and analyze data as described in our recent methods article (Berger and Chistol, 2024). Below we summarize the data analysis pipeline specific for the single molecule experiments analyzed in this study.

**Selecting DNA Molecules for Downstream Analysis**

Regions of interest (ROIs) corresponding to individual DNA molecules were selected manually based on the previews of DNA stained with SYTOX Orange (these were acquired before the start of the experiment). Only well-separated DNA molecules were used in the analysis.

**Analyzing Polymerase Binding/Dissociation Events**

To analyze the binding and dissociation of polymerases we employed an approach previously described in (Terui et al., 2024) for analyzing replication initiation factors. ROIs corresponding to polymerase binding events were selected from edges of FEN1 tracts, corresponding to the position of the replisome.

To analyze the binding and dissociation of polymerases on singly tethered molecules, ROIs were also selected from the DNA Preview – singly tethered molecules appear as diffraction limited sources of SYTOX signal. Only singly-tethered molecules that replicated (displayed FEN1 signal) were analyzed. To minimize the number of "non-productive" binding events (Figure S1E) analyzed we applied the following criteria: total FEN1 signal was integrated over time. This signal increases as the molecule replicates but eventually peaks and decreases presumably due to completion of DNA replication triggering FEN1 unloading. Polymerase binding events were therefore only selected from time points prior to the peak in FEN1 signal.

**Changepoint Analysis**

To identify the time when lagging strand synthesis ceased following chase with a catalytically dead mutant (Figure 3D,G) the edge of the FEN1 tract was fit with the following bilinear model in MATLAB:

$$b(x) = \begin{cases} a_1 x + b & x < x_0 \\ a_2 x + (a_1 - a_2) b x_0 & x \geq x_0 \end{cases}$$

where $x_0$ is the changepoint time, $a_1$ is the slope before the changepoint, $a_2$ is the slope after the changepoint, and $b$ is the y-intercept. To confirm the robustness of the changepoint analysis we compared the goodness of fit to a linear model that lacks a changepoint term. The following monlinear model was fit to the same FEN1 tract:

$$m(x) = a_1 x + b$$

For each model we approximated the Bayesian Information Criterion (BIC) of the fit using:

$$BIC \approx n \log\left(\frac{sse}{n}\right) + K\,log(n)$$

where $n$ is the sample size, $sse$ is the sum of squared errors of the fit, and $K$ is the number of parameters in the model (Cavanaugh and Neath, 2019). We computed the difference of the BIC scores for each FEN1 tract:

$$\Delta BIC = BIC_{monolinear} - BIC_{bilinear}$$

where larger ΔBIC values indicate the FEN1 tract was better fit by the bilinear model. Comparing the ΔBIC values across the mock, Pol $\alpha^{Dead}$, and Pol $\delta^{Dead}$ chases confirmed that the mutant chases were better modeled by the bilinear fitting than the mock chase (Figure S3A-B). This approach yields a changepoint for the control chase even though the FEN1 tract does not actually stall. To confirm that the changepoint identified in the control is statistically distinct from the mutant chase experiments we performed pairwise 2-sample Kolmogorov Smirnov tests for the three conditions.

**CMG Velocity Analysis**

To measure CMG velocity during/after the chase we fit the GINS signal with a linear model in MATLAB and extracted the slope. Only timepoints that occurred after the chase were used for fitting.
